# Food perception alone stimulates oviposition through serotonin signaling in *Drosophila*

**DOI:** 10.64898/2026.10.08.757633

**Authors:** Tze Chang Ng, Madison Oh, Ayush Shrivastava, Christian Arntsen, Sai Sankpal, Matthew W. Chong, Advaitha Kamalakkannan, Jinhu Chung, Molly Stanley, Yang Lyu

## Abstract

Reproduction is strongly shaped by nutrient availability, but whether sensory information about food can regulate reproductive output before nutrients are consumed remains unclear. Here, using *Drosophila melanogaster*, we show that sensory perception of dietary yeast, a protein-rich food source widely used to stimulate fecundity in the fruit fly, is sufficient to rapidly promote egg laying. Specifically, we found that yeast odor exposure increased the rate of oviposition and the number of eggs laid within hours, independent of yeast intake, and that this response was mediated by olfactory signaling through the odorant receptor co-receptor Orco. We further identified serotonin receptor 2A (5-HT2A), but not 5-HT2B, as necessary for yeast odor-induced egg laying. We then asked whether yeast odor also elicits broader life-history changes and whether its reproductive and longevity effects depend on a shared mechanism. Consistent with previous observations, yeast odor also shortened lifespan. However, this lifespan effect was independent of 5-HT2A signaling, indicating that the reproductive and longevity responses to food perception are separable. Finally, our brain connectomic analyses revealed that yeast-responsive olfactory neurons preferentially influence oviposition descending neurons (OviDNs) compared with other olfactory neurons. OviDNs further connect with endocrine-related circuitry in the corpora allata (CA), and, most interestingly, the interneurons linking OviDNs to CA neurons are enriched for serotonergic and dopaminergic modulation. Taken together, our observations indicate that nutrient acquisition is not a prerequisite for animals to adjust their reproduction to the nutritional environment. Sensory information predicting food availability itself can elicit changes in neural state, likely through olfactory-reproductive circuits, that acutely alter reproductive physiology.

**Significance statement:** Food perception can drive changes beyond feeding-related behaviors. This study establishes that olfactory perception of dietary yeast per se elicits egg-laying behavior in *Drosophila melanogaster* and that the conserved neuromodulator serotonin is required for this response. Our observations suggest that the reproductive state of animals is determined not only by commonly recognized factors such as ingested nutrients, but also by sensory cues that anticipate food availability.

## Introduction

For many species, including humans, reproduction is strongly influenced by nutrient availability (1–4). In the fruit fly, *Drosophila melanogaster*, dietary protein promotes oviposition and oocyte development and influences reproductive senescence (5–7). This dietary influence is perhaps not surprising, given the substantial nutritional demands of reproduction (8–10) and the role of amino acid-sensing pathways in linking nutrient availability to reproductive output (11–13). Most studies of dietary protein and reproduction, however, have focused on nutrients after they are consumed within the animal. More recently, studies in model systems have shown that the sensory perception of food itself can influence physiology and lifespan (14–18). In *Caenorhabditis elegans* and *D. melanogaster*, specific sensory circuits that detect the odor or taste of food modulate aging in response to perceived nutrient availability (14, 15, 17, 19–21). These observations suggest that reproductive responses may be initiated by food perception even before the nutrients that support reproduction are consumed.

At the molecular level, sensory information is integrated through neuromodulation to coordinate behavioral and physiological outcomes (22–25). Serotonin signaling regulates feeding, metabolism, and lifespan in response to perceived food abundance and nutrient availability (14, 17, 19–21). Work in *Drosophila* and *C. elegans* has shown that serotonergic neurons interact with taste, olfactory, and mechanosensory circuits to modulate responses to food-related cues (24, 26, 27). In *C. elegans*, serotonin signaling also coordinates germline proliferation and egg laying in response to sensory cues from male pheromones (28, 29). Consequently, we speculated that serotonin signaling encodes external food availability through sensory circuits to influence reproductive decisions.

Using *D. melanogaster*, we tested the hypothesis that food perception per se influences reproductive and long-term life-history decisions independently of nutrient intake. We removed dietary yeast from the fly chamber while simultaneously exposing flies to yeast odor (Fig. 1A), allowing us to isolate the effects of yeast perception from the effects of direct contact with dietary yeast. Indeed, our results show that yeast odor exposure alone increased egg laying within a few hours, likely through the detection of yeast-derived volatiles. Consistently, we found that this response required olfactory signaling through the odorant receptor co-receptor Orco. More importantly, yeast odor-induced oviposition also required serotonin signaling, with a specific role for serotonin receptor 2A (5-HT2A), but not serotonin receptor 2B (5-HT2B). Although yeast odor also shortened lifespan, as observed in previous studies (14, 15, 17), this effect was surprisingly independent of 5-HT2A signaling. These results indicate that the reproductive and lifespan responses to perceived food availability can be decoupled and mediated through distinct molecular pathways. Bulk RNA-seq and connectomic analyses suggest that olfactory neural circuits may encode perceived nutrient availability and coordinate reproductive and other life-history responses. Together, our findings highlight that reproductive state is influenced not only by the nutrients an animal consumes, but also by sensory cues that anticipate food availability.

**Figure 1.**
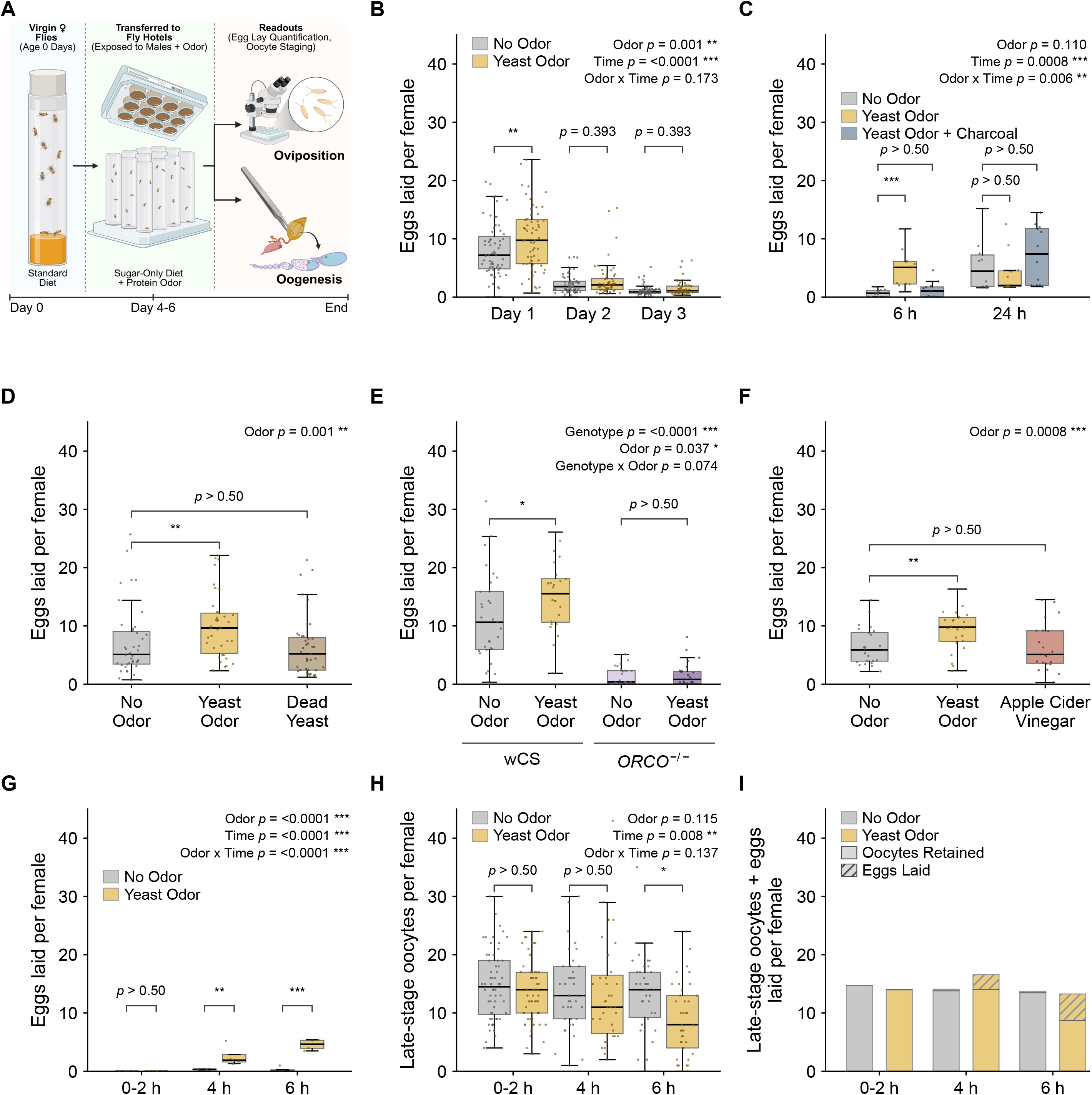
Yeast odor stimulates oviposition in *D. melanogaster* through olfactory detection. **(A)** Experimental setup. Virgin females were collected on eclosion, aged, and housed with males in either a chamber (“fly hotel”) of 12 individual vials (4 × 3), or a standard fly cage (not shown). Each vial or cage contained 10 females and 10 males. Egg laying and ovary morphology were measured as readouts of oviposition and oogenesis, respectively. **(B)** Yeast odor increases egg laying over three consecutive days (n = 62–63 vials per group, 4 experimental blocks). **(C)** Activated charcoal suppresses the effects of yeast odor on egg laying at 6 h, but not at 24 h (n = 10–12 vials per group, 2 experimental blocks). **(D)** Microwaved, dead yeast fails to stimulate egg laying (n = 38–42 vials per group, 4 experimental blocks). **(E)** Loss of *Orco* abolishes the egg–laying response to yeast odor. *Orco*^−/−^ constitutive mutants were compared with their background control, wCS (n = 24–31 vials per group, 3 experimental blocks). **(F)** Apple cider vinegar, an odorant attractive to *Drosophila*, does not reproduce the effect of live yeast odor (n = 23–24 vials per group, 2 experimental blocks). **(G)** Yeast odor rapidly increases egg laying. Eggs laid per female were counted at 0, 1, 2, 4, and 6 h of odor exposure (n = 4–6 cages per group, 3 experimental blocks). **(H)** Yeast odor depletes the pool of mature oocytes. Late-stage (S13–S14) oocytes were counted in females dissected from cages in (G) at the respective time points (n = 38–60 females per group, 3 experimental blocks). **(I)** Yeast odor shifts reproductive output from retaining oocytes to laying eggs. Eggs laid (G) and late-stage (S13–S14) oocytes retained (H) were summed per female and stacked (n = 4–6 cages per group, 3 experimental blocks). Boxes represent the median and interquartile range, with whiskers extending to 1.5× IQR. Asterisks denote \**p* < 0.05, \*\**p* < 0.01, and \*\*\**p* < 0.001; non-significant comparisons are labeled with their exact *p*-value, or as “*p* > 0.50” where *p* exceeds 0.50. Full statistical details are in Table S1.

## Results

### Yeast odor rapidly stimulates egg laying in the absence of dietary protein

Reproduction requires substantial nutrient investment, and female fecundity in *Drosophila* is strongly stimulated by dietary protein (6, 9, 30). However, previous studies have shown that food perception per se can influence longevity and survival (14–17, 31), raising the possibility that perceived food availability may also influence reproductive decisions. Because dietary yeast is a major protein source for flies and strongly promotes female fecundity (9, 32), we exposed adult females to yeast odor while removing yeast from the food medium (Fig. 1A). Yeast odor increased egg laying compared with flies not exposed to yeast odor, with the strongest effect observed on the first day of exposure (Fig. 1B and Supplementary Table 1, ANOVA, *p* = 0.001). Thus, yeast odor alone is sufficient to promote oviposition in the absence of dietary protein intake.

To determine whether this response depends on volatile cues produced by live yeast, we first removed yeast-derived odorants with charcoal. This treatment initially abolished the increase in egg laying, although the effect of charcoal diminished over time (Fig. 1C). Second, microwaved yeast failed to increase egg laying (Fig. 1D, Tukey HSD post hoc, *p* > 0.50), further supporting a requirement for volatile cues produced by live yeast. We next examined flies lacking the odorant receptor co-receptor Orco, which is required for the function of the canonical odorant receptor family in *Drosophila* (33, 34). Loss of Orco abolished the yeast odor-induced increase in egg laying (Fig. 1E, Tukey HSD post hoc, *p* > 0.50), demonstrating that olfactory signaling is required for this response. Apple cider vinegar, a strong attractant for fruit flies (35), did not increase egg laying (Fig. 1F, Tukey HSD post hoc, *p* > 0.50), indicating that attraction to an odor source alone is insufficient. Together, these results show that volatile cues produced by live yeast elicit oviposition through the olfactory sensory system.

Given the increase in egg laying during yeast odor exposure, we sought to distinguish whether yeast odor only stimulates egg release or also promotes egg production and maturation. We focused on stage 13 and 14 oocytes, which are mature or near-mature oocytes available for ovulation and subsequent oviposition (36, 37). Interestingly, females began laying eggs 4 h after mating regardless of yeast odor exposure, whereas yeast odor increased the number of eggs laid over this period, with the difference becoming larger by 6 h (Fig. 1G). Despite the increased oviposition, yeast-exposed females retained slightly fewer late-stage oocytes at 6 h (Fig. 1H). To further determine whether yeast odor exposure increases total reproductive output, we combined the number of eggs laid with the number of late-stage oocytes retained in the ovary. This total measure was similar between yeast odor-exposed and control females (Fig. 1H). Overall, although yeast odor rapidly promotes egg laying, we did not detect a noticeable increase in the total number of late-stage oocytes and eggs laid. These results indicate that the acute effect of yeast odor is mainly on oviposition rather than oogenesis.

### 5-HT2A, not 5-HT2B, modulates yeast odor-induced egg laying

Serotonin has been implicated in sensing internal protein status and establishing protein intake targets in *Drosophila* (19, 21), as well as in mediating physiological responses to external food availability in both *Drosophila* and *C. elegans* (14, 20, 38). We therefore speculated that serotonin signaling might link perceived nutrient availability to reproductive decisions. We first tested the role of 5-HT2A, a type-2 excitatory G protein-coupled serotonin receptor, as we and others have previously found that loss of 5-HT2A alters protein preference and protein homeostasis in *Drosophila* (19–21). Additionally, we studied the involvement of 5-HT2B, a second type-2 serotonin receptor (39), to understand whether the potential effects are specific to individual receptors or reflect a broader role for type-2 serotonin signaling.

We first examined egg laying in constitutive 5-HT2A and 5-HT2B loss-of-function mutants and their genetic background control (w; Canton-S). Control females began laying eggs approximately 4-6 h after mating, and yeast odor strongly increased egg laying by 6 h (Fig. 2A; Tukey HSD, *p* < 0.0001; Supplementary Table 2). Both *5-HT2A* and *5-HT2B* mutants laid fewer eggs than control females. Loss of 5-HT2A completely abolished the increase in egg laying in response to yeast odor (Tukey HSD, *p* > 0.5), whereas *5-HT2B* mutants retained a modest response that did not reach statistical significance (Tukey HSD, *p* = 0.083). Together, although both receptors contribute to normal egg laying, our data indicate that 5-HT2A is specifically required for the acute ovipositional response to yeast odor.

**Figure 2.**
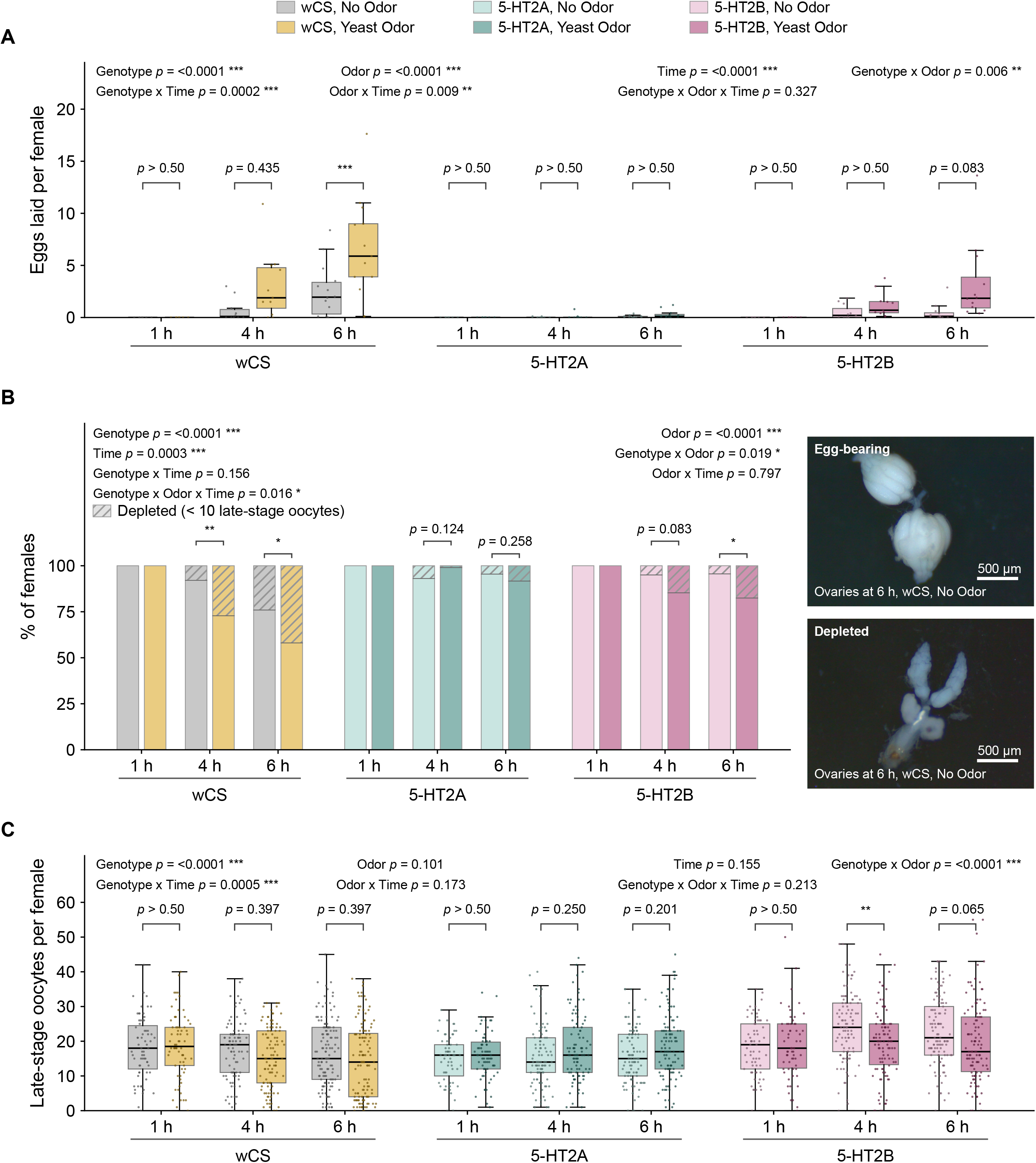
Serotonin receptor 5-HT2A, but not 5-HT2B, is specifically required for the oviposition response to yeast odor. **(A)** Loss of either 5-HT2A or 5-HT2B reduces egg laying. Yeast odor increases egg laying in control (wCS) flies by 6 h, but this increase is abolished in 5-HT2A mutants and reduced in 5-HT2B mutants (n = 7–14 cages per group, 5 experimental blocks). **(B)** Left: yeast odor increases the proportion of oocyte-depleted ovaries in control (wCS) flies. This response is lost in 5-HT2A mutants but retained in 5-HT2B mutants (n = 62–133 females per group, 5 experimental blocks). Ovaries were scored as oocyte-depleted if they contained fewer than 10 late-stage (S13–S14) oocytes and came from a cage in which eggs were laid. Females scored at 1 h were excluded from statistical analysis as no eggs were laid in any genotype at that time point. Right: representative oocyte-bearing and oocyte-depleted ovaries (wCS, no odor, 6 h; scale bars, 500 µm). **(C)** Yeast odor does not significantly alter the number of late-stage (S13–14) oocytes at any time point in control or 5-HT2A mutant flies, but decreases it in 5-HT2B mutants at 4 h (n = 62–133 females per group, 5 experimental blocks). Boxes represent the median and interquartile range, with whiskers extending to 1.5x IQR. Asterisks denote \**p* < 0.05, \*\**p* < 0.01, and \*\*\**p* < 0.001; non-significant comparisons are labeled with their exact *p*-value, or as “*p* > 0.50” where *p* exceeds 0.50. Full statistical details are in Table S2.

Although we observed reduced egg laying in serotonin receptor mutants, this impairment could result either from reduced oviposition or from a decrease in the pool of mature oocytes available to be laid. To distinguish between these possibilities, we examined late-stage (S13/S14) oocytes. We first quantified the proportion of ovaries depleted of late-stage oocytes (Fig. 2B). Consistent with the egg-laying response, yeast odor increased the proportion of depleted ovaries at 4 and 6 h after exposure (Tukey HSD, *p* = 0.006 and *p* = 0.014, respectively). Loss of 5-HT2A abolished this yeast odor-induced increase in ovarian depletion (Tukey HSD, *p* = 0.124 at 4 h and *p* = 0.258 at 6 h). In contrast, 5-HT2B mutants still exhibited the depletion response, with a trend at 4 h and a significant effect by 6 h (Tukey HSD, *p* = 0.083 and *p* = 0.014, respectively). These results further support a specific requirement for 5-HT2A in the ovipositional response to yeast odor.

We next quantified the number of late-stage oocytes in the ovary and found that, across genotypes and time points, yeast odor did not significantly alter late-stage oocyte abundance overall (Fig. 2C and Supplementary Table 2; linear mixed-effects model, odor effect, *p* = 0.101). We observed a notable difference in 5-HT2B mutants, in which yeast odor reduced the number of late-stage oocytes relative to unexposed mutants. This trend was similar to that observed in wild-type flies, suggesting that yeast odor increased egg laying and ovarian depletion at the same time (Fig. 2A and B). Together with the results shown in Fig. 1, our analysis of these mutants further indicates that yeast odor acutely elicits oviposition rather than increasing the pool of late-stage oocytes and, critically, that this response is specifically mediated by 5-HT2A signaling.

### Reproductive and lifespan responses to yeast odor are separable

Previous studies in both *D. melanogaster* (15) and *C. elegans* (14, 17) have shown that exposure to food odors can partially suppress the longevity benefits of dietary restriction and reduce lifespan. Because yeast odor also acutely promotes oviposition, we asked whether its reproductive and longevity effects reflect a shared life-history response and whether 5-HT2A signaling contributes to both.

Consistent with previous findings (15), exposure to yeast odor shortened the lifespan of control females maintained on a restricted SY5 diet (Fig. 3A; log-rank test, *p* = 0.01). We next asked whether 5-HT2A, which is required for yeast odor-induced oviposition, also mediates the effect of yeast odor on lifespan. In the absence of yeast odor, *5-HT2A* mutants showed a modest increase in lifespan relative to their *w*; CS background controls (Fig. 3A; log-rank test, *p* = 0.01). However, loss of 5-HT2A did not abolish the lifespan-shortening effect of yeast odor (Fig. 3A, log-rank test, *p* < 0.001), in contrast to its requirement for the ovipositional response (Figs. 2A and B). In other words, yeast odor continued to shorten lifespan in *5-HT2A* mutants despite their failure to increase oviposition in response to odor. Thus, the acute ovipositional and lifespan responses to perceived food availability can be genetically separated, with 5-HT2A required for the former but not the latter.

**Figure 3.**
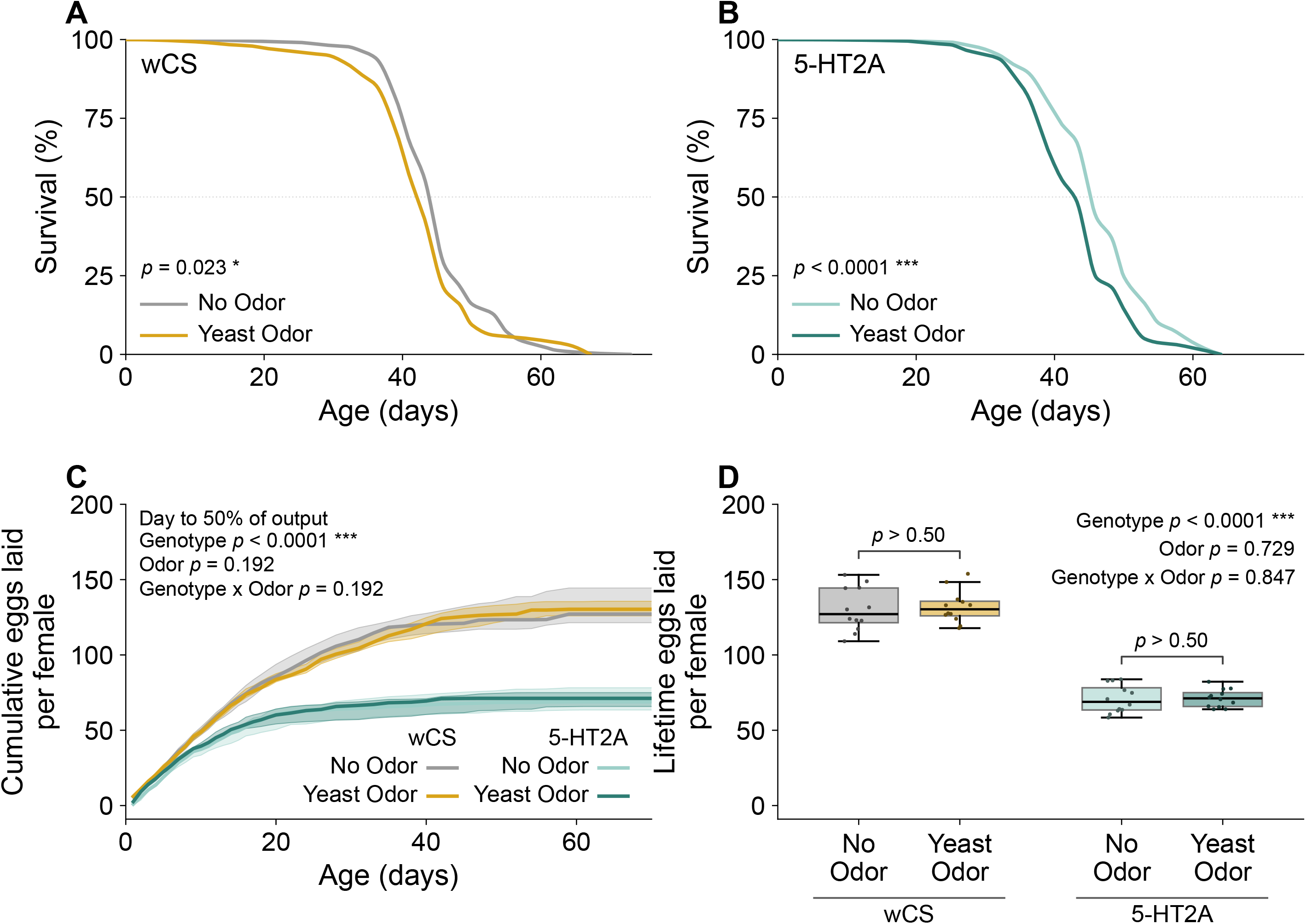
Yeast odor shortens lifespan independently of 5-HT2A, without altering lifetime fecundity. **(A, B)** Constant yeast odor shortens the lifespan in both control (wCS) flies (A) and 5-HT2A mutants (B). Females were maintained on SY5 medium with or without continuous yeast odor and scored until death (n = 243-245 females per group, 20 per vial at setup). Curves show Kaplan-Meier survival estimates; survival was compared using a log-rank test. **(C)** Yeast odor does not alter the accumulation of eggs over adult life in either genotype, whereas 5-HT2A mutants lay fewer eggs and plateau earlier (n = 12 vials per group, 15 per vial at setup). Lines show the group median, and shaded bands indicate the interquartile range. The day on which each vial reached 50% of its total output was analyzed using two-way ANOVA for genotype, odor, and their interaction, with terms tested by Type II sums of squares. **(D)** Yeast odor does not alter lifetime egg production in either genotype at day 70 (endpoint of C). Lifetime eggs per female were compared by the same two-way ANOVA, followed by Tukey HSD across the four groups; the two within-genotype odor comparisons are shown. Boxes represent the median and interquartile range, with whiskers extending to 1.5x IQR. Asterisks denote \**p* < 0.05, \*\**p* < 0.01, and \*\*\**p* < 0.001; non-significant comparisons are labeled with their exact *p*-value, or as “*p* > 0.50” where *p* exceeds 0.50.

Because acute ovipositional behavior may not fully reflect cumulative reproductive output over the lifespan, we further examined the effects of yeast odor and 5-HT2A signaling on lifetime fecundity. In contrast to its acute effect on egg laying, yeast odor did not alter lifetime fecundity (Figs. 3B and 3C; two-way ANOVA, odor effect, *p* = 0.729), suggesting that the reproductive effect of food odor reflects acute regulation of oviposition rather than an increase in cumulative reproductive investment. Loss of 5-HT2A, however, substantially reduced lifetime fecundity (Figs. 3B and 3C; two-way ANOVA, genotype effect, *p* < 0.001), consistent with its reduced egg laying at earlier time points (Fig. 2A). Yeast odor did not alter lifetime fecundity in *5-HT2A* mutants (Figs. 3B and 3C; two-way ANOVA, genotype × odor interaction, *p* = 0.847), despite its effects on their lifespan (Fig. 3A). Together, these results indicate that the lifespan-shortening effect of yeast odor does not require 5-HT2A and is uncoupled from its 5-HT2A-dependent acute effects on oviposition. Moreover, yeast odor did not alter lifetime fecundity, further arguing against a simple reproduction-longevity trade-off as the basis for the lifespan effect.

### Loss of 5-HT2A signaling broadly alters transcription, whereas yeast odor has limited transcriptomic effects

To determine how food perception and 5-HT2A signaling influence the head and ovary, we analyzed transcriptomes from control and 5-HT2A mutant females exposed to either yeast odor or no odor for 4 h. Principal component analysis (PCA) revealed that tissue identity accounted for the largest source of variation, with head and ovary samples separating along PC1 (87.7%; Fig. 4A). Genotype also contributed to the variation, with control and 5-HT2A mutant samples separating along PC3 (1.2%; Fig. 4A). Notably, genotype-dependent separation along PC3 was more pronounced in head samples than in ovaries, consistent with the enriched expression of 5-HT2A in the nervous system (40).

**Figure 4.**
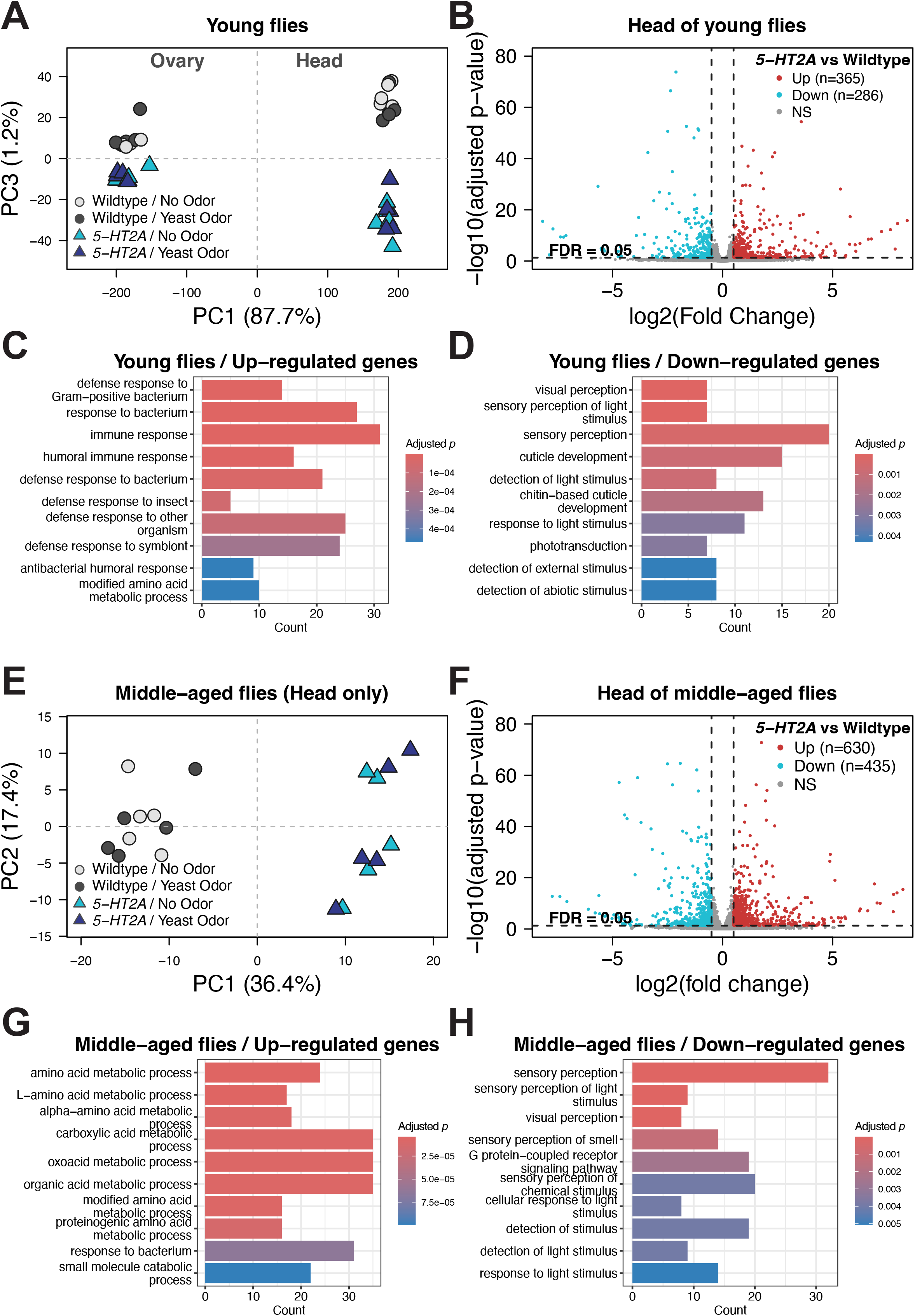
Bulk RNA-seq reveals age-specific and tissue-specific changes induced by loss of 5-HT2A signaling. **(A, E)** Principal component analysis (PCA) shows the distribution of samples from young flies (A; head and ovary) and middle-aged flies (E; head only). In young flies, transcripts contributing strongly to PC1 and PC3 distinguish tissues and genotypes. In the heads of middle-aged flies, transcripts contributing strongly to PC1 separate the genotypes. n = 10 samples per group, with ∼40 heads or ∼20 ovaries used for each sample. **(B, F)** Volcano plots show differentially expressed genes (DEGs) between 5-HT2A mutants and control (wCS) flies in the heads of young (B) and middle-aged (F) flies. DEGs were defined using FDR ≤ 0.05 and |log2FC| ≥ 0.5. Roughly equal numbers of up- and down-regulated genes were observed at each age, while middle-aged flies exhibited a greater total number of DEGs than young flies. **(C, D, G, H)** Gene Ontology (GO) analysis showing the top 10 categories enriched among up- or down-regulated genes in young and middle–aged flies. Biological Process (BP) terms are shown here; all significantly enriched categories (adjusted P ≤ 0.05) for BP, Molecular Function (MF), and Cellular Component (CC) are provided in Supplementary Table 3. Categories enriched among down–regulated genes were similar between young and middle-aged flies, whereas up-regulated genes were enriched in different categories, suggesting age–specific effects.

We first examined the transcriptional effects associated with loss of 5-HT2A. Using a linear mixed model, we identified 651 differentially expressed genes in the head and 295 in the ovary between control and 5-HT2A mutant females (FDR ≤ 0.05, |log2FC| ≥ 0.5; Fig. 4B and Supplementary Fig. 1). In the head, all five enriched Gene Ontology (GO) Biological Process (BP) terms among genes upregulated in 5-HT2A mutants were related to immune processes (Fig. 4C and Table 2). Notably, four of the five enriched BP terms among downregulated genes were related to sensory processes, including olfactory and visual responses (Fig. 4D and Table 2). Additional Cellular Component (CC) and Molecular Function (MF) terms were enriched among differentially expressed genes in the head of 5-HT2A mutants (Supplementary Table 3). In the ovary, no significant GO terms were identified among neither up- nor down-regulated genes.

We then asked whether the rapid reproductive response to yeast odor is associated with broader changes in transcript levels. Despite its strong effect on oviposition within hours, 4 h of yeast odor exposure resulted in only three differentially expressed genes (DEGs) in the head (MFS14, CG13059, and CG3348) and none in the ovary (FDR ≤ 0.05, |log2FC| ≥ 0.5). We speculated that prolonged exposure might produce more extensive changes in expression profiles and therefore examined 20-day-old females maintained on SY5 medium with or without yeast odor throughout adult life. However, PCA again showed little separation between the yeast odor-exposed and unexposed groups (Fig. 4E). These results suggest that yeast odor exposure have relatively little effect on global patterns of gene expression in the tissues examined.

Loss of 5-HT2A signaling led to widespread changes in transcript levels in middle-aged flies (Fig. 4F). To explore whether these changes are age-specific, we performed GO analysis on DEGs (FDR ≤ 0.05, |log2FC| ≥ 0.5) identified in the head of middle-aged 5-HT2A mutants. Among the upregulated genes, we found that enriched terms were dominated by metabolic processes (Fig. 4G and Supplementary Table 3). This pattern differed from the profile of upregulated genes in younger mutant flies, where enriched GO terms were largely related to immune function. Furthermore, genes downregulated in middle-aged 5-HT2A mutants were enriched for sensory processes, including olfactory and visual responses, similar to those observed in younger 5-HT2A mutants (Fig. 4H and Supplementary Table 3). Our bulk RNA-seq results indicate that the transcriptional effects of 5-HT2A vary with age, whereas yeast odor itself leads to little detectable alteration in global expression profiles. It is possible that the effects of yeast odor are confined to a relatively small population of neurons or involve mechanisms that do not require widespread changes in transcript levels.

### Yeast-responsive olfactory circuits influence oviposition and reproductive endocrine pathways

We next sought to understand how yeast-sensing olfactory neurons may communicate with neural circuits associated with egg laying and reproduction. The recently completed Female Adult Brain and Nerve Cord (BANC) connectome (41) provides an opportunity to trace olfactory inputs to circuits associated with these distinct reproductive processes. We focused on two potential targets. Oviposition descending neurons (OviDNs) control the oviposition motor program and integrate information about egg-laying substrate value to regulate when and where eggs are deposited (42, 43). We also examined the corpora allata (CA), the major source of juvenile hormone (JH), given its key role in regulating female reproduction in insects (44, 45). We therefore asked whether yeast-responsive olfactory receptor neurons (ORNs) connect to OviDNs and the CA through distinct neural pathways.

Using a proofread subset of the BANC connectome (v888), we identified 535 olfactory receptor neurons (ORNs) that respond to yeast-associated compounds, hereafter referred to as yeast-responsive ORNs (Fig. 5A and Supplementary Video). Based on community-provided annotations, 460 reside in the antenna and 75 in the maxillary palp. Shortest-path analysis showed that all but one of the yeast-responsive ORNs were connected to 16 oviposition descending neurons (OviDNs) within five synaptic hops. Since signals may traverse multiple pathways simultaneously, we also used a linear dynamical model of signal transduction developed by Bates et al. (41) to compute the influence of source neurons on target neurons. This analysis further showed that yeast-responsive ORNs exerted greater influence on OviDNs than other ORNs (Supplementary Fig. 2). When we compared their influence on OviDNs with that on all other descending neurons (DNs; n = 1,301), yeast-responsive ORNs showed a clear enrichment for OviDNs (Fig. 5B). Moreover, they showed the strongest OviDN selectivity among ORN subclasses (Fig. 5B).

**Figure 5.**
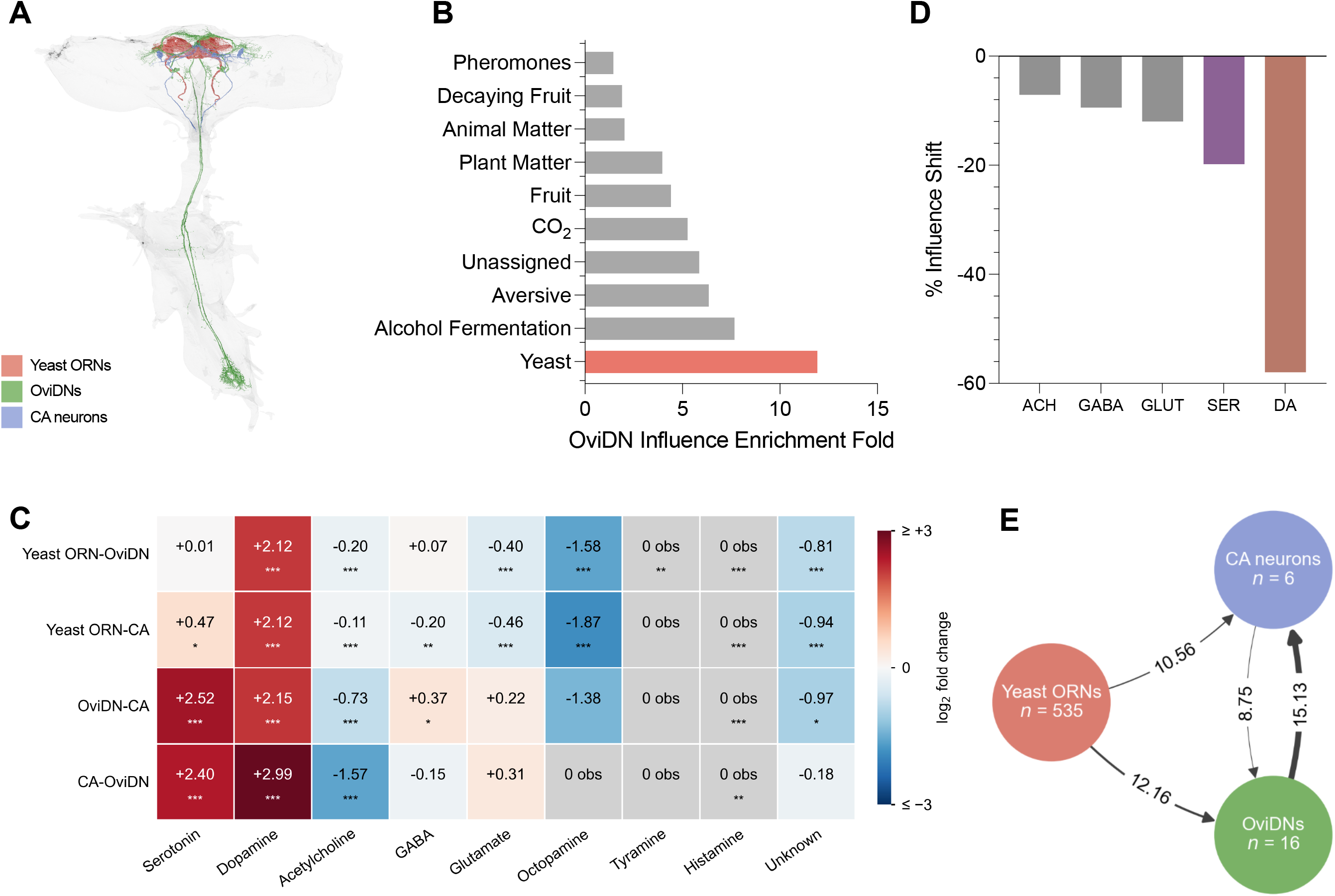
Yeast-responsive olfactory receptor neurons (ORNs) preferentially influence oviposition descending neurons (OviDNs), which reach the *corpura allata* (CA)-projecting neurons through serotonergic and dopaminergic interneurons. **(A)** Morphology of yeast-responsive ORNs (yeast ORNs; red), OviDNs (green), and CA-projecting neurons (blue) in the BANC connectome (n = 535 yeast ORNs, 16 OviDNs, 6 CA-projecting neurons). Yeast ORNs are defined as olfactory receptor neurons functionally tuned to fermentation cues, annotated as “yeasty_volatile” in the BANC connectome. **(B)** Yeast ORNs show the strongest OviDN selectivity of any ORN subclass. For each ORN, influence enrichment was calculated as its mean influence on OviDNs (n = 16) divided by its mean influence on all other descending neurons (n = 1,301), then averaged within its subclass (e.g., Yeast, Alcohol Fermentation). Influence was computed with the linear dynamical model of Bates et al. 2026. The reported scores are fold changes of raw influence scores. The corresponding comparison between yeast ORNs and all other ORNs is in Supplementary Fig. 2. **(C)** Neurons along the shortest OviDN-CA and CA-OviDN circuits are predicted to be enriched for serotonin and dopamine. Neurotransmitter enrichment in yeast ORN-OviDN, yeast ORN-CA, OviDN-CA, and CA-OviDN circuits was tested against connectome-wide proportions using a two-sided Fisher’s exact test with Benjamini-Hochberg correction. Asterisks denote \**p* < 0.05, \*\**p* < 0.01, and \*\*\**p* < 0.001; cells without asterisks were not significant after correction. “0 obs” indicates that the circuit contained no neuron of that neurotransmitter; where such a cell carries an asterisk, it denotes significantly fewer neurons than expected, rather than a fold change. **(D)** Silencing serotonergic (SER) and dopaminergic (DA) interneurons reduces OviDN to CA influence more than silencing cholinergic (ACH), GABAergic, or glutamatergic (GLUT) interneurons. The OviDN to CA-projecting neuron circuit was reconstructed across four layers (Supplementary Fig. 4), and its second-order interneurons were stratified by neurotransmitter and silenced *in silico*. The reported changes in influence derive from raw influence scores. **(E)** A large population of yeast ORNs (n = 535) influences smaller populations of OviDN (n = 16) and CA-projecting neurons (n = 6), with the strongest influence running from OviDN to CA-projecting neurons. Values are influence scores (Bates et al. 2026), log-transformed, where 1 unit is a 2.7-fold difference in influence. Arrow width is proportional to the score.

In contrast, we detected only weak influence between yeast-responsive ORNs and six CA neurons (Supplementary Fig. 3A). However, the OviDNs showed strong and specific connectivity with these CA neurons (Supplementary Fig. 3B). These connectivity patterns suggest that yeast-responsive olfactory information can influence endocrine pathways involved in oogenesis indirectly through OviDN-to-CA connectivity.

Lastly, we characterized the neuronal populations contributing to these connections and the involvement of serotonin. Neurons along the shortest OviDN-CA pathways were enriched for both serotonin and dopamine (Fig. 5C). To determine which interneurons were the most influential in this circuit, we reconstructed three- and four-layer OviDN-CA pathways and identified interneurons that are directly postsynaptic to OviDNs (Supplementary Fig. 4). This gave us 37 neurons of five neurotransmitter types as candidates for silencing. We then recalculated OviDN-CA influence after silencing each neuronal population in silico. Our analysis indicated that silencing serotonergic neurons reduced circuit influence more than silencing cholinergic, GABAergic, or glutamatergic neurons (Fig. 5D), revealing a modulatory role for serotonin in this circuitry. To our surprise, silencing dopaminergic neurons caused an even larger reduction, indicating that dopamine also contributes strongly to this pathway alongside serotonin. Taken together, our observations suggest a potential circuit architecture through which olfactory information and neuromodulatory signaling may coordinate multiple reproductive outputs (Fig. 5E).

## Discussion

For decades, *Drosophila* biologists have used yeast paste on egg-laying media to stimulate oviposition and collect large numbers of progeny, and this effect has generally been thought to be due to the nutritional value of ingested yeast. However, yeast-derived volatiles provide strong olfactory cues. Becher et al. argued that yeast-derived volatiles contribute to attraction and oviposition in *D. melanogaster*, although the sensory and nutritional components of yeast were not experimentally isolated in that study (46). More recently, several studies have shown that olfactory and gustatory cues modulate oviposition-site preference in *Drosophila* (47–50), and that these egg-laying choices differ across species in ways that reflect their ecological niches. For instance, in the noni specialist *D. sechellia*, olfactory input is important for oviposition on its host fruit, although the same study demonstrated that noni odor alone was not sufficient to increase egg laying (49), in contrast to our finding that yeast odor alone rapidly stimulates oviposition in *D. melanogaster*. A different study showed that ripe-strawberry volatiles promote oviposition on ripe fruit in the ripe-fruit specialist *D. suzukii* but reduce oviposition in *D. melanogaster* (50). These observations illustrate that sensory information plays a critical role in determining where females lay their eggs. However, whether food perception alone can influence whether and how quickly females reproduce before nutrients are consumed has not been rigorously tested.

In this study, by exposing flies to yeast odor without allowing direct contact with dietary yeast, we experimentally isolated the effects of perceived yeast availability and probed the underlying mechanisms. Strikingly, we found that yeast odor alone was sufficient to trigger a rapid increase in egg laying through an olfactory–5-HT2A signaling axis. Quantification of late-stage oocytes further indicated that yeast odor solely increased oviposition without increasing the number of late-stage oocytes. In other words, the acute reproductive response to yeast odor acts largely on egg laying rather than on the production of additional mature oocytes. In nature, yeast-derived odor may serve as a long-range cue for females to locate a nearby oviposition site. Anticipatory food-associated sensory cues may therefore be sufficient to shift females toward a neural state tied to subsequent reproductive physiology.

At the neural level, how food-associated sensory signals influence downstream neural circuits remains much less understood. Dweck et al. showed that activation of Or71a-expressing olfactory neurons promotes both egg laying and feeding (48), whereas Guillemin et al. more recently found that activation of Ir94e-expressing gustatory neurons promotes egg laying while mildly suppressing feeding (12). Leveraging the power of published connectomic data, we discovered strong connections between food-specific olfactory neurons and the oviposition circuit. More specifically, yeast-responsive ORNs showed particularly strong influence on OviDNs compared with other descending neurons, providing a potential circuit mechanism through which yeast-derived olfactory information influences oviposition. Interestingly, our influence analysis further showed that the crosstalk between OviDNs and CA neurons is strongly impacted by serotonergic and dopaminergic neurons. Overall, we believe that the underlying circuits are complex, and that interneurons linking sensory neurons and reproductive circuits form neuromodulatory networks that may allow these signals to produce nuanced behavioral and physiological responses. Notably, the different effects of Or71a- and Ir94e-expressing neurons on feeding (12, 48) suggest that sensory regulation of egg laying does not simply reflect a general increase in food-related behaviors.

Our genetic analysis further identifies 5-HT2A as a key component linking food perception to reproductive output. Compared with the well-established roles of serotonin in feeding (19, 21, 24, 51), metabolism (20, 51, 52), and longevity (14, 19, 20, 53, 54), how serotonin regulates reproductive investment remains less clear. We previously found that 5-HT2A signaling in *Drosophila* promotes an energetically costly metabolic state in response to the nutritional environment (20). Here, we further show that loss of 5-HT2A abolished the increase in egg laying induced by yeast odor, providing a novel link between food perception and reproductive output. In parallel, our bulk RNA-seq data indicate that 5-HT2A signaling is associated with broader systemic changes involving trade-offs among distinct physiological programs. Specifically, sensory processes were enriched among genes reduced by loss of 5-HT2A, whereas immune-related processes were enriched among genes increased in young 5-HT2A mutants, and metabolic processes, including amino acid metabolism, were enriched among genes increased in middle-aged 5-HT2A mutants. We therefore propose that 5-HT2A has a systems-level role in coordinating sensory, metabolic, immune, and reproductive responses to the nutritional environment. Such an integrative role for serotonin may be broadly conserved, as studies in *C. elegans* have shown that serotonin signaling coordinates germline proliferation and egg laying and regulates germline investment in response to environmental cues (28, 29).

Our study also echoes the long-standing idea from life-history theory that reproduction carries costs for survival (55–57). Previous studies in long-lived *Drosophila* have associated delayed oocyte maturation (58) and altered yolk-related physiology (59) with increased lifespan. Our observations indicate that life-history decisions can be influenced not only by nutrients that an animal consumes, but also by the perceived availability of those nutrients. In natural environments, animals regularly rely on sensory information to make behavioral and physiological decisions. Specific cues associated with food, mates, and danger can provide immediate indications about environmental conditions before direct interaction occurs. In the case of mating, prior work has demonstrated that perceiving pheromones from the opposite sex is sufficient to alter physiology and lifespan even in the absence of mating (60–62). Here, we extend this framework to food perception by showing that yeast odor alone is sufficient to stimulate egg laying before nutrients are consumed. Consistent with previous studies (14, 15), we also found that yeast odor shortened lifespan in addition to its short-term effect on reproduction. However, the lifespan effect was clearly independent of 5-HT2A signaling, indicating that the reproductive and longevity effects of food perception are mediated through separable mechanisms. Overall, our study suggests that animals may use sensory cues to anticipate resource availability and calibrate reproductive physiology before nutrient acquisition, while short- and long-term life-history responses to food perception are mediated through different molecular pathways.

## Materials and Methods

The fly strains used in this study are described below. *D. melanogaster* w[*]; Canton-S (wCS) and *Orco*^-/-^ flies (originally from the Leslie Vosshall laboratory) were obtained from the Scott Pletcher laboratory at the University of Michigan. *5-HT2A*^[PL00052]^ mutants (BDSC #19367) and *5-HT2B*^[GAL4]^ mutants (BDSC #86278) were originally obtained from the Bloomington *Drosophila* Stock Center and backcrossed to wCS for 7-10 generations.

### Fly husbandry and maintenance

All fly stocks were maintained on a standard cornmeal-based larval growth medium (produced by LabScientific Inc and purchased from Fisher Scientific) and in a controlled environment (25 °C, 60% humidity) with a 12:12 hr light:dark cycle. We controlled the developmental larval density by manually aliquoting 32 µl of collected eggs into individual bottles containing 25 ml of food. Immediately after eclosion, virgin female flies were collected via anesthetization using CO2 briefly and allowed to age for 4-6 days. Remaining mixed-sex flies were kept on SY10 (10% [w/v] sucrose and 10% [w/v] yeast) medium for 2-3 days until they were used for experiments. Sucrose (purchased from Labviva) and MP Biomedicals Brewer’s Yeast (purchased from Fisher Scientific) were used in our study.

### Yeast exposure and oviposition quantification

All assays used wCS flies unless specified otherwise. Virgin female flies were housed in either 5.0 cm x 7.3 cm (diameter x height) chambers (10 flies per chamber) or in fly hotels consisting of 12 individual vials (Fig. 1A) and maintained on a sucrose only diet (10% [w/v] sucrose). Flies were exposed to yeast odor for the duration specified for each experiment, with matched cohorts maintained without yeast odor as controls. Yeast odor was generated by a thick yeast paste prepared from 20g Fleischman’s Instant Dry Yeast and 30mL distilled water. The yeast paste was placed approximately 1 inch above the mesh of the chamber or fly hotel to allow volatiles to reach the flies without direct contact. Experimental chambers and odor sources were housed within airtight containers to create a closed environment that allowed odorants to diffuse while limiting exposure to external odors. Containers were stored in incubators at 25°C and (65-85% humidity) under a 12-hour light/dark cycle.

Female flies were acclimated to their respective experimental conditions for 24 h before 3-6-day-old males were introduced at a 1:1 ratio to allow mating. Containers were opened only for data collection or for replacement of fresh food and odor-producing reagents every 24 h. Egg laying was quantified at the indicated time points.

For additional experiments, the standard yeast paste was modified or replaced to test the contribution of specific odor properties. For charcoal adsorption experiments, 5 g of activated charcoal (Nutricost) was mixed with the yeast paste to reduce yeast-derived volatile cues. To compare live and heat-killed yeast, yeast paste was microwaved at high power for 2 min, or until a pungent fermentative odor developed, before exposure to flies. To determine whether the oviposition response reflected general attraction to an odor source, we tested apple cider vinegar, a known attractant for *Drosophila*. Two beakers containing 20 mL of apple cider vinegar (365 by Whole Foods Market) were placed in each closed container, allowing volatile cues to reach the flies by diffusion.

### Late-stage oocyte quantification

Flies were exposed to either no-odor or yeast-odor conditions as described above. Reproductive output was measured at 2, 4, and 6 h by quantifying the number of eggs laid per fly and dissecting ovaries to determine the total number of late-stage (stage 13/14; S13/S14) oocytes, as described previously (63). For oocyte quantification, flies were dissected in 70% ethanol.

### Lifespan and life-time fecundity measurements

Two-day-old mated female flies were sorted into fly hotels at 15 flies per vial and exposed to either yeast odor or no odor as the control condition. Each fly hotel was housed in a separate container to prevent odor transfer between conditions. Flies were maintained on a homogeneous SY5 diet containing 5% (w/v) sucrose and yeast at 25°C and 65-85% relative humidity under a 12 h light/12 h dark cycle. Humidity was additionally controlled by placing two small Petri dishes, each containing 10 g of silica gel beads (Dry & Dry), inside each container, as continuous release of volatiles from the yeast paste contributed to excessive condensation. Food and yeast paste were replaced daily. Survival was quantified by daily population census, and fecundity was measured by manually counting the number of eggs laid on each food plate throughout the lifespan.

### Total RNA extraction and DNase removal

Experimental flies were briefly submerged in 70% ethanol then dissected in PBS to collect both heads and ovaries (n=40 heads or 20 ovaries per sample). Dissected tissue was immediately transferred to 1.7 ml microcentrifuge tubes containing 200 μl of ice-cold TRIzol reagent and homogenized with an RNase-free pestle. An additional 800 μl ice-cold TRIzol was added, and samples were processed by vigorous shaking with 200 μl of chloroform. The resulting aqueous phase was subject to isopropanol precipitation with GlycoBlue for pellet visibility. RNA pellets were washed twice with ice-cold 75% ethanol, air-dried, and resuspended in nuclease-free water. RNA concentration and purity were assessed using NanoDrop.

For DNase treatment, 5 μl of 10x Incubation Buffer and 1 μl DNase were added to each sample on ice. After brief incubation at room temperature for 20 minutes, samples were returned to ice and mixed with 500μl isopropanol by gentle inversions. After a 10-min incubation at room temperature to precipitate RNA, pellets were washed with ice-cold 75% ethanol as described above, air dried, and resuspended in 50 μl of IDTE pH 7.5 (10 mM Tris, 0.1 mM EDTA). Samples were kept on ice before a final concentration check.

RNA concentrations were then diluted to an acceptable concentration (100 ng/μl). RNA integrity was verified through gel electrophoresis and TapeStation RNA ScreenTape assay. Approved samples were submitted at 50 ng/μl to Plasmidsaurus for RNA-Seq (3′ end counting, Illumina NovaSeq X Plus).

### Analysis of high-throughput RNA sequencing

Single-end sequencing reads were trimmed to remove adapter sequences and low-quality base reads using FastP (64, 65). We obtained ∼19.3M reads for each sample (18.2-20.3M; n=60 libraries across two developmental cohorts), with a mean read length of 91.3 bp. Preprocessed reads were aligned to the *Drosophila melanogaster* transcriptome (The GTF annotation file is obtained from ftp.flybase.net/genomes/Drosophila_melanogaster/, version: r6.68), using STAR (66) (parameters: -runThreadN 4 -runMode genome Generate), with gene annotations obtained from FlyBase (release r6.68; ftp.flybase.net/genomes/Drosophila_melanogaster/). Gene-level read counts were calculated using featureCounts (67) by counting reads overlapping annotated exons. Differential gene expression analysis was performed using DeSEQ2 (68), which models gene-level counts using a negative binomial generalized linear model. The model included genotype, treatment, and their interaction (∼ genotype + treatment + genotype:treatment). Genes with a Benjamini–Hochberg adjusted *p*-value (false discovery rate [FDR]) < 0.05 were considered differentially expressed. Gene Ontology (GO) (69) enrichment analysis was performed using the R packages clusterProfiler (70), AnnotationDbi, and org.Dm.eg.db.

### Connectomic analysis

All Adjusted Influence analyses were conducted in Python (v3.13.13 and v3.14.3) using the connectome Influence Calculator from the Drugowitsch Lab (v2.0.0, https://github.com/DrugowitschLab/ConnectomeInfluenceCalculator) developed for the BANC connectome (41). Based on the Drugowitsch Lab’s implementation, custom code was adapted to run the Influence Calculator using modifiable parameters and to process the calculator’s raw output (https://github.com/ayush-shrivastava003/banc-analysis). The BANC connectome (v888) was analyzed using the corresponding connectivity (banc_888_edgelist_simple_v3.feather) and metadata (banc_888_meta.feather) files that were downloaded from the Harvard dataverse (https://dataverse.harvard.edu/dataset.xhtml?persistentId=doi:10.7910/DVN/7WTH1N). Synaptic connections containing fewer than 3 synapses were excluded and the remaining edges were normalized to the postsynaptic neuron’s total input (syn_weight_measure = “norm”). Additionally, the full connectivity matrix was rescaled to a maximum eigenvalue of λ = 0.99. Based on recommendations from Bates et al., the metadata table was restricted to neurons that were annotated as “proofread” or “roughly_proofread”. Because of this filtering, some neurons from each FlyWire-derived population were excluded from influence analysis. The exact neuron IDs used for influence analysis can be found on our GitHub (https://github.com/ayush-shrivastava003/banc-analysis). When calculating influence from a sensory neuron or sensory population, all non-seed neurons annotated as “sensory” or “ascending_sensory” were computationally silenced. Computational silencing prevents specified neurons from propagating influence through the network (https://github.com/DrugowitschLab/ConnectomeInfluenceCalculator), thereby preventing the recruitment of additional sensory pathways and restricting propagation to independent pathways originating from the seed population.

For calculating the population-level influence scores between yeast ORNs, OviDNs, and CA neurons (Fig. 5B, S5A, S5B), all seed neurons were activated simultaneously so that the resulting influence reflects the combined activity of the full source population. This approach generates a combined raw influence score for each target neuron. Each target’s raw influence was then divided by the number of seed neurons, and these normalized values were averaged across target neurons to obtain a single population-level influence value. Consistent with the analysis used by Bates et al., raw population-level influence scores were converted to “Adjusted Influence” using a log-transformation and a constant shift of c = 24. To calculate the individual neuron-level influence scores and influence enrichment between ORNs and DNs/OviDNs (Fig. 5B), each seed ORN was activated individually so that the resulting influence reflects the activity of just that single source neuron. This approach generates an all-seeds x all-targets raw influence score table. For each ORN seed, its mean raw influence was calculated by averaging across all the target neuron scores. Specifically, each seed’s mean influence to OviDNs (n=16) and each seed’s mean influence to all other (non-OviDN) descending neurons (DNs) (n=1301) were computed. Within a given ORN seed, OviDN enrichment was calculated (mean raw OviDN influence/mean raw DN influence) This process was repeated for each ORN population. Because enrichment was defined as a fold ratio between influence values, this analysis was performed using raw influence rather than log-transformed Adjusted Influence scores.

For influence analyses combined with neuronal population silencing (Fig. 5D), population-level influence between OviDNs and CA neurons was calculated using simultaneous seed activation to establish a baseline level of circuit influence. Candidate second-order neurons in this circuit were identified using a 4-layer circuit construction pipeline. The code used to map this circuit and identify the putative second-order populations can be found on our Github. Briefly, neurons immediately post-synaptic to OviDNs (second-order/Layer 2) and neurons immediately pre-synaptic to CA neurons (Layer 3) were identified using a minimum synapse threshold of 3. Only second-order neurons that connected to either to Layer 3 neurons or CA neurons directly were retained. Among these second-order circuit neurons, there were 2 acetylcholine (ACH), 9 dopamine (DA), 17 GABA, 3 glutamate (GLUT), and 6 serotonin (SER) neurons. To determine the relative impact each population has on overall OviDN-CA influence, each population was computationally silenced (see above) and circuit influence was re-calculated. The impact of each population was quantified by measuring the percent loss of OviDN-CA circuit influence following silencing relative to the intact-network baseline (%Influence Shift). Raw influence scores, not log-transformed Adjusted Influence values, were used to calculate %Influence Shift.

For Fig. 5D, we first determined the shortest unweighted pathways between each population using breadth-first searches of the connectome. Next, for each circuit, we identified the number of neurons affiliated with each neurotransmitter type listed in the metadata file: serotonin, dopamine, acetylcholine, GABA, glutamate, octopamine, tyrosine, and histamine. Those without a neurotransmitter type listed were marked “unknown”. We then determined the concentration of these neurotransmitter types in the entire connectome. For each neurotransmitter in each circuit, we calculated the log2(fold change) and performed a two-sided Fisher’s exact test against the connectome-wide concentrations. We applied Benjamini-Hochberg correction and only considered those with aN FDR < 0.05 significant.

Fig. 5A and Supplementary Video were rendered using Blender and Python. The code for this and all other analyses are available on GitHub at github.com/ayush-shrivastava003/banc-analysis for connectomic analysis.

## Data and code availability

RNA-seq data have been deposited in GEO under accession GSE349230 and are publicly available as of the date of publication. Microscopy data reported in this paper will be shared by the corresponding author upon request. All original code has been deposited on GitHub at github.com/ylyu-fly/yeast-odor-rnaseq for RNA-seq analysis and at github.com/ayush-shrivastava003/banc-analysis for connectomic analysis. Any additional information required to reanalyze the data reported in this paper is available from the lead contact upon request.

## Supporting information

Table S1. Statistical analyses for Figure 1

Table S2. Statistical analyses for Figure 2

Table S3. GO term enrichment summary, related to Figure 4

Supplementary Video. 3D rendering of yeast-responsive ORNs, OviDNs, and CA-projecting neurons in the BANC connectome, related to Figure 5A

## Acknowledgments

The authors thank members of the Lyu laboratory for assistance with fly husbandry, reagent preparation, and genetic backcrossing, as well as our departmental colleagues for their generous support in establishing the laboratory. We also thank the Kim McKim laboratory at Rutgers University for guidance and expertise in fly ovary dissection and stage 13/14 oocyte quantification.

This work was supported by Rutgers start-up funds to Y.L. C.A. and M.S. were supported by research funds from the University of Vermont. This research was conducted while Yang Lyu was a Glenn Foundation for Medical Research and AFAR Grant for Junior Faculty awardee. M.O. was supported by a Rutgers Division of Life Sciences Summer Undergraduate Research Fellowship (DLS-SURF). Research reported in this publication was supported by the National Institute on Aging of the National Institutes of Health under grant number R01AG095099 to Y.L. Ten percent of this research was financed with federal funds, and 90% was financed with non-governmental funds.

## Disclaimer

The content is solely the responsibility of the authors and does not necessarily represent the official views of the National Institutes of Health.

---

**Supplementary Fig. 1.**
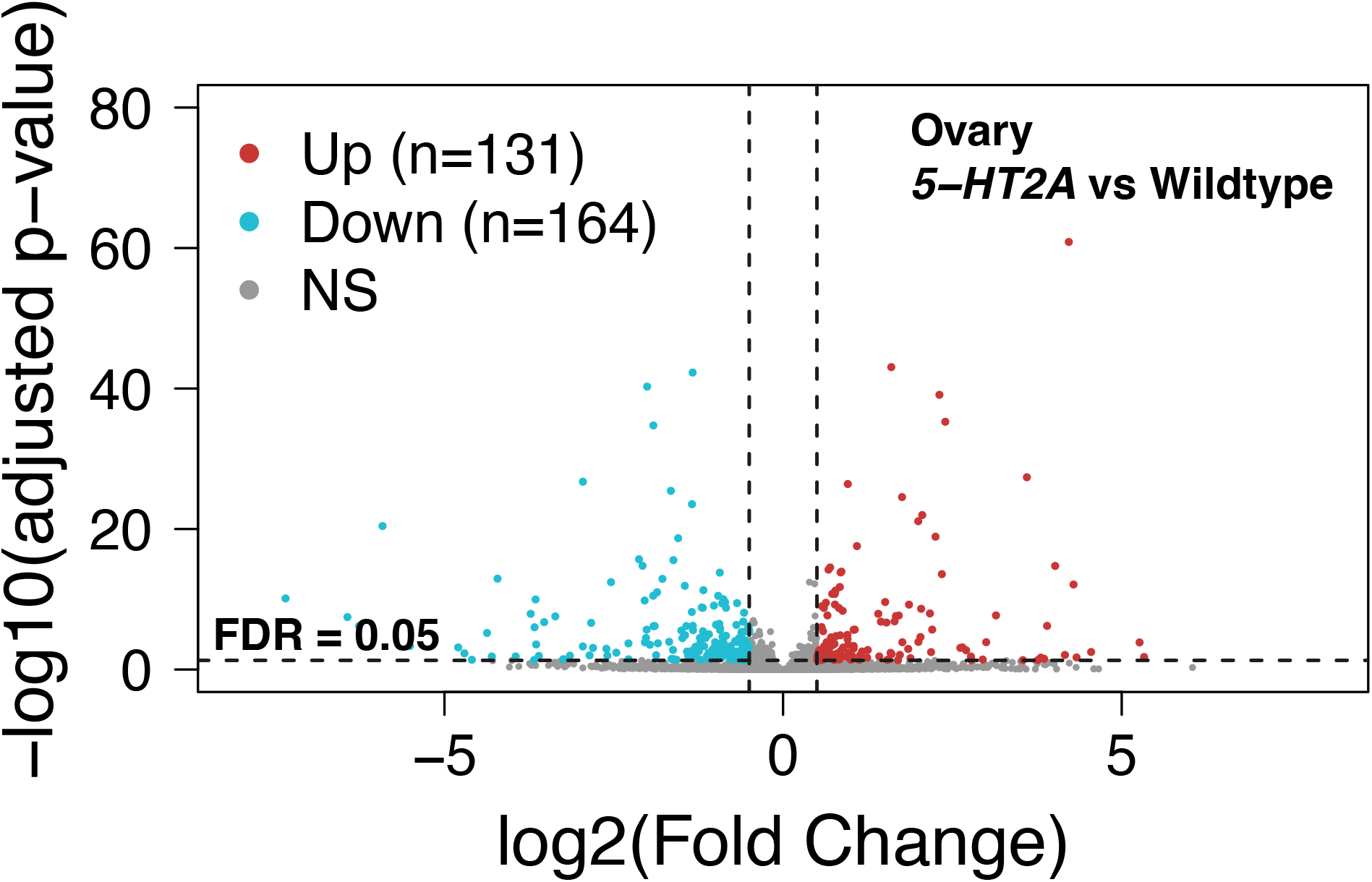
Volcano plots show differentially expressed genes (DEGs) between 5-HT2A mutants and control (wCS) flies in the ovaries of young flies.

**Supplementary Fig. 2.**
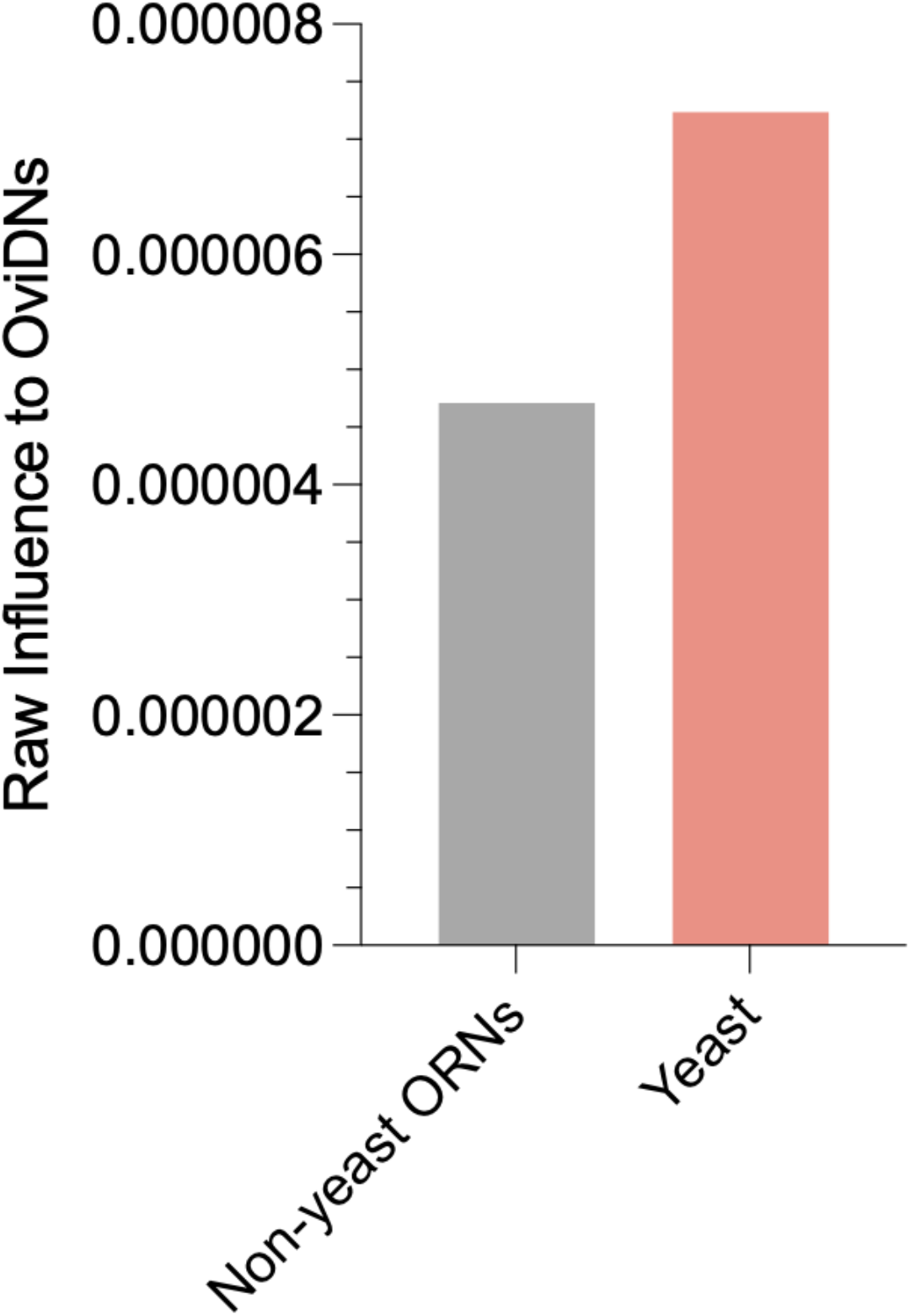
OviDNs are influenced more strongly by yeast-responsive ORNs over other ORNs. Population-level raw influence on OviDNs (n = 16) from yeast-responsive ORNs (n = 535) or from all other ORNs. Influence was summed across target neurons and normalized by the number of seed neurons, yielding a single value per group; no error bars or statistical test apply to a population-level score.

**Supplementary Fig. 3.**
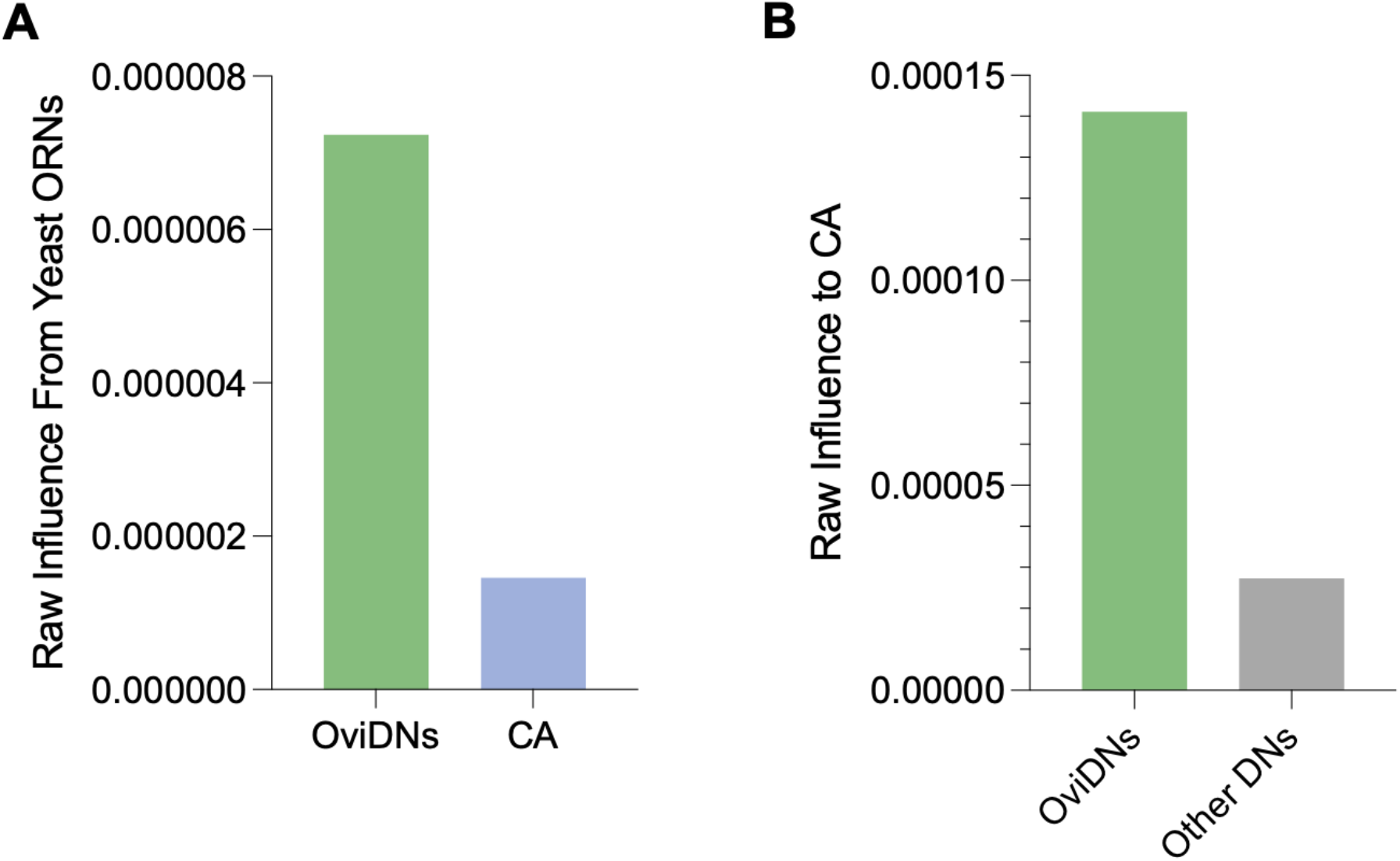
CA-projecting neurons are influenced more strongly by OviDNs than yeast-responsive ORNs or by other descending neurons (DNs). Related to Fig 5. **(A)** Population-level raw influence of yeast-responsive ORNs (n = 535) on OviDNs (n = 16) or on CA-projecting neurons (n = 6). **(B)** Population-level raw influence of OviDNs (n = 16) or of all other descending neurons (n = 1,301) on CA-projecting neurons (n = 6). Each bar represents a single population-level value, so no error bars or statistical tests apply.

**Supplementary Fig. 4.**
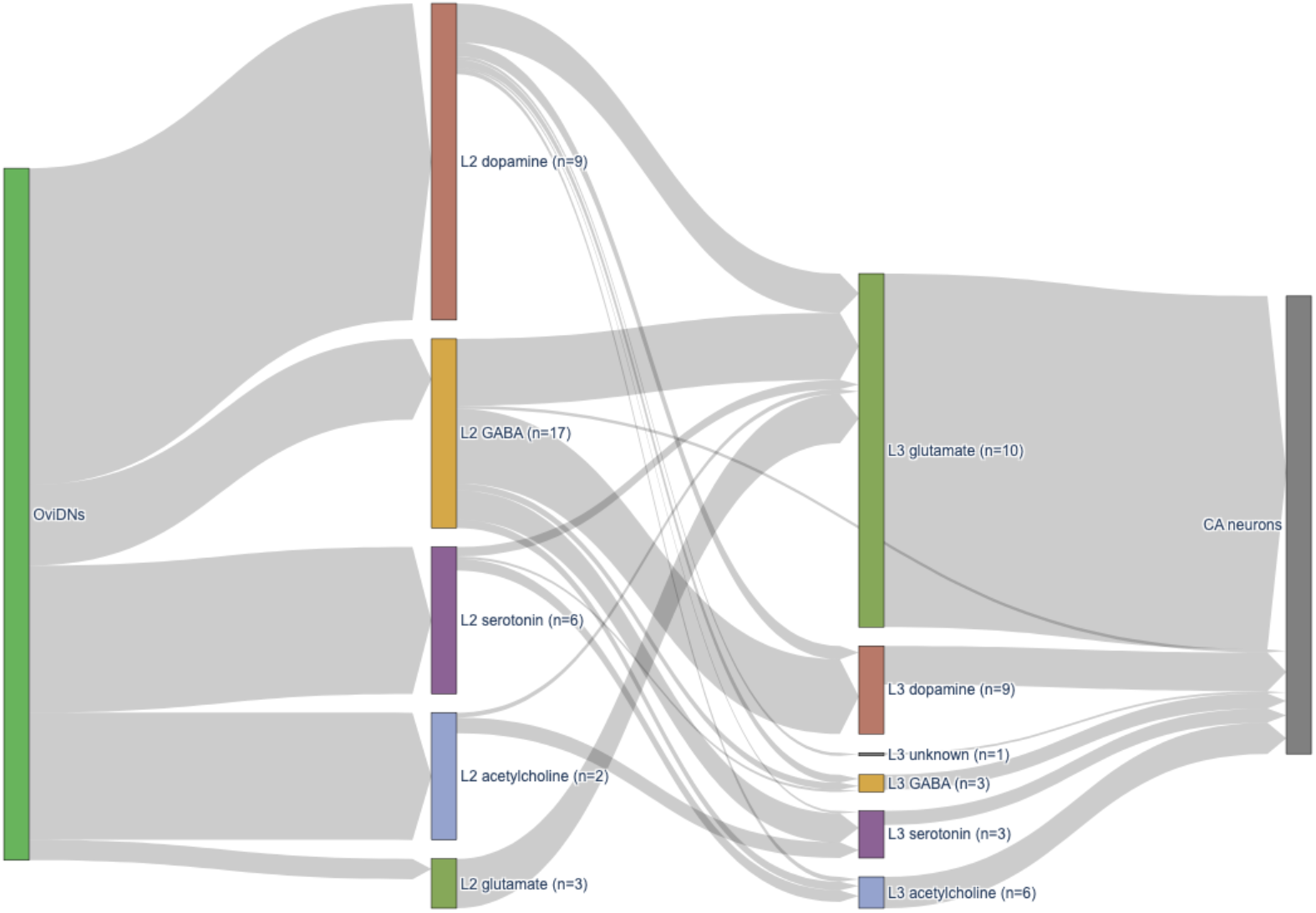
Reconstruction of the four-layer OviDN to CA-projecting neuron circuit. Related to Fig 5. Flow diagram of the neurons connecting OviDNs (left) to CA-projecting neurons (right), grouped by layer and by neurotransmitter. Each band’s width is proportional to the total number of incoming and outgoing synapses for that group of neurons. L2 = layer 2 (second order); L3 = layer 3 (third order). The L2 populations (2 acetylcholine, 9 dopamine, 17 GABA, 3 glutamate, and 6 serotonin neurons) were the candidates silenced in Figure 5D; L3 populations (6 acetylcholine, 9 dopamine, 3 GABA, 10 glutamate, 3 serotonin, and 1 unknown) are shown for completeness.

## References

1. J. P. Moatt, S. Nakagawa, M. Lagisz, C. A. Walling, The effect of dietary restriction on reproduction: a meta-analytic perspective. BMC Evol Biol 16, 199 (2016).

2. N. M. Templeman, C. T. Murphy, Regulation of reproduction and longevity by nutrient-sensing pathways. J Cell Biol 217, 93–106 (2018).

3. T. Iwasa et al., Effects of low energy availability on female reproductive function. Reprod Med Biol 21, e12414 (2022).

4. A. B. Loucks, Energy availability, not body fatness, regulates reproductive function in women. Exerc Sport Sci Rev 31, 144–148 (2003).

5. C. K. Mirth, A. Nogueira Alves, M. D. Piper, Turning food into eggs: insights from nutritional biology and developmental physiology of *Drosophila*. Curr Opin Insect Sci 31, 49–57 (2019).

6. D. A. Skorupa, A. Dervisefendic, J. Zwiener, S. D. Pletcher, Dietary composition specifies consumption, obesity, and lifespan in *Drosophila melanogaster*. Aging Cell 7, 478–490 (2008).

7. B. G. Ruchitha et al., Effect of developmental and adult diet composition on reproductive aging in *Drosophila melanogaster*. Exp Gerontol 194, 112501 (2024).

8. D. Drummond-Barbosa, A. C. Spradling, Stem cells and their progeny respond to nutritional changes during *Drosophila* oogenesis. Dev Biol 231, 265–278 (2001).

9. T. J. Bradley, F. H. Simmons, An analysis of resource allocation in response to dietary yeast in *Drosophila melanogaster*. J Insect Physiol 43, 779–788 (1997).

10. E. Bowman, M. Tatar, Reproduction regulates *Drosophila* nutrient intake through independent effects of egg production and sex peptide: Implications for aging. Nutr Healthy Aging 4, 55–61 (2016).

11. A. R. Armstrong, K. M. Laws, D. Drummond-Barbosa, Adipocyte amino acid sensing controls adult germline stem cell number via the amino acid response pathway and independently of Target of Rapamycin signaling in *Drosophila*. Development 141, 4479–4488 (2014).

12. J. Guillemin et al., Taste cells expressing Ionotropic Receptor 94e reciprocally impact feeding and egg laying in *Drosophila*. Cell Rep 43, 114625 (2024).

13. L. LaFever, A. Feoktistov, H. J. Hsu, D. Drummond-Barbosa, Specific roles of Target of rapamycin in the control of stem cells and their progeny in the *Drosophila* ovary. Development 137, 2117–2126 (2010).

14. H. A. Miller et al., Serotonin and dopamine modulate aging in response to food odor and availability. Nat Commun 13, 3271 (2022).

15. S. Libert et al., Regulation of *Drosophila* life span by olfaction and food-derived odors. Science 315, 1133–1137 (2007).

16. Y. Luo et al., Yeast volatiles double starvation survival in *Drosophila*. Sci Adv 7 (2021).

17. B. Zhang, H. Jun, J. Wu, J. Liu, X. Z. S. Xu, Olfactory perception of food abundance regulates dietary restriction-mediated longevity via a brain-to-gut signal. Nat Aging 1, 255–268 (2021).

18. C. E. Riera et al., The sense of smell impacts metabolic health and obesity. Cell Metab 26, 198–211.e195 (2017).

19. J. Ro et al., Serotonin signaling mediates protein valuation and aging. Elife 5 (2016).

20. Y. Lyu et al., *Drosophila* serotonin 2A receptor signaling coordinates central metabolic processes to modulate aging in response to nutrient choice. Elife 10 (2021).

21. A. S. Munneke, T. S. Chakraborty, S. S. Porter, C. M. Gendron, S. D. Pletcher, The serotonin receptor 5-HT2A modulates lifespan and protein feeding in *Drosophila melanogaster*. Front Aging 3, 1068455 (2022).

22. Z. Q. Tang, L. O. Trussell, Serotonergic modulation of sensory representation in a central multisensory circuit Is pathway specific. Cell Rep 20, 1844–1854 (2017).

23. Z. Li et al., Dissecting a central flip-flop circuit that integrates contradictory sensory cues in *C. elegans* feeding regulation. Nat Commun 3, 776 (2012).

24. Z. Yao, K. Scott, Serotonergic neurons translate taste detection into internal nutrient regulation. Neuron 110, 1036–1050.e1037 (2022).

25. E. S. Kitto, S. Beydoun, S. F. Leiser, Rewarding touch limits lifespan through neural to intestinal signaling. Proc Natl Acad Sci U S A 122, e2423780122 (2025).

26. X. Zhang, Q. Gaudry, Functional integration of a serotonergic neuron in the *Drosophila* antennal lobe. Elife 5 (2016).

27. A. Schoofs et al., Serotonergic modulation of swallowing in a complete fly vagus nerve connectome. Curr Biol 34, 4495–4512.e4496 (2024).

28. E. Z. Aprison, S. Dzitoyeva, I. Ruvinsky, The serotonin circuit that coordinates germline proliferation and egg laying with other reproductive functions in *Caenorhabditis elegans*. Proc Biol Sci 289, 20220913 (2022).

29. E. Z. Aprison, S. Dzitoyeva, I. Ruvinsky, The roles of TGFβ and serotonin signaling in regulation of germline progenitors in adult *C. elegans* hermaphrodites revealed by the effects of a male pheromone. Dev Biol 534, 183–195 (2026).

30. K. Jensen, C. McClure, N. K. Priest, J. Hunt, Sex-specific effects of protein and carbohydrate intake on reproduction but not lifespan in *Drosophila melanogaster*. Aging Cell 14, 605–615 (2015).

31. J. Apfeld, C. Kenyon, Regulation of lifespan by sensory perception in *Caenorhabditis elegans*. Nature 402, 804–809 (1999).

32. H. Colinet, D. Renault, Dietary live yeast alters metabolic profiles, protein biosynthesis and thermal stress tolerance of *Drosophila melanogaster*. Comp Biochem Physiol A Mol Integr Physiol 170, 6–14 (2014).

33. M. C. Larsson et al., Or83b encodes a broadly expressed odorant receptor essential for *Drosophila* olfaction. Neuron 43, 703–714 (2004).

34. R. Benton, S. Sachse, S. W. Michnick, L. B. Vosshall, Atypical membrane topology and heteromeric function of *Drosophila* odorant receptors in vivo. PLoS Biol 4, e20 (2006).

35. J. L. Semmelhack, J. W. Wang, Select *Drosophila* glomeruli mediate innate olfactory attraction and aversion. Nature 459, 218–223 (2009).

36. C. Berg, M. Sieber, J. Sun, Finishing the egg. Genetics 226 (2024).

37. A. C. Spradling, Developmental genetics of oogenesis. The development of Drosophila melanogaster, 1–70 (1993).

38. E. V. Entchev et al., A gene-expression-based neural code for food abundance that modulates lifespan. Elife 4, e06259 (2015).

39. F. Saudou, U. Boschert, N. Amlaiky, J. L. Plassat, R. Hen, A family of *Drosophila* serotonin receptors with distinct intracellular signalling properties and expression patterns. Embo j 11, 7–17 (1992).

40. C. D. Nichols, 5-HT2 receptors in *Drosophila* are expressed in the brain and modulate aspects of circadian behaviors. Dev Neurobiol 67, 752–763 (2007).

41. A. S. Bates et al., Distributed control circuits across a brain-and-cord connectome. Nature 10.1038/s41586-026-10735-w (2026).

42. F. Wang et al., Neural circuitry linking mating and egg laying in *Drosophila* females. Nature 579, 101–105 (2020).

43. V. Vijayan et al., A rise-to-threshold process for a relative-value decision. Nature 619, 563–571 (2023).

44. R. Yamamoto, H. Bai, A. G. Dolezal, G. Amdam, M. Tatar, Juvenile hormone regulation of *Drosophila* aging. BMC Biol 11, 85 (2013).

45. C. G. Santos, F. C. Humann, K. Hartfelder, Juvenile hormone signaling in insect oogenesis. Curr Opin Insect Sci 31, 43–48 (2019).

46. P. G. Becher et al., Yeast, not fruit volatiles mediate *Drosophila melanogaster* attraction, oviposition and development. Funct Ecol 26, 822–828 (2012).

47. H. K. Dweck et al., Olfactory preference for egg laying on citrus substrates in *Drosophila*. Curr Biol 23, 2472–2480 (2013).

48. H. K. Dweck, S. A. Ebrahim, A. Farhan, B. S. Hansson, M. C. Stensmyr, Olfactory proxy detection of dietary antioxidants in *Drosophila*. Curr Biol 25, 455–466 (2015).

49. R. Álvarez-Ocaña et al., Odor-regulated oviposition behavior in an ecological specialist. Nat Commun 14, 3041 (2023).

50. S. Lebreton et al., An Orco dependent pathway suppresses egg-laying attraction to ripe fruit volatile compounds in *Drosophila melanogaster* but not in *D. suzukii*. BMC Ecol Evol 25, 102 (2025).

51. S. Srinivasan et al., Serotonin regulates *C. elegans* fat and feeding through independent molecular mechanisms. Cell Metab 7, 533–544 (2008).

52. J. Luo, J. Becnel, C. D. Nichols, D. R. Nässel, Insulin-producing cells in the brain of adult *Drosophila* are regulated by the serotonin 5-HT1A receptor. Cell Mol Life Sci 69, 471–484 (2012).

53. T. S. Chakraborty et al., Sensory perception of dead conspecifics induces aversive cues and modulates lifespan through serotonin in *Drosophila*. Nat Commun 10, 2365 (2019).

54. S. F. Leiser et al., Cell nonautonomous activation of flavin-containing monooxygenase promotes longevity and health span. Science 350, 1375–1378 (2015).

55. T. Flatt, Survival costs of reproduction in *Drosophila*. Exp Gerontol 46, 369–375 (2011).

56. T. B. Kirkwood, Evolution of ageing. Nature 270, 301–304 (1977).

57. G. C. Williams, Natural selection, the costs of reproduction, and a refinement of Lack’s principle. The American Naturalist 100, 687–690 (1966).

58. K. Carlson, T. Nusbaum, M. Rose, L. Harshman, Oocyte maturation and ovariole number in lines of *Drosophila melanogaster* selected for postponed senescence. Functional Ecology, 514–520 (1998).

59. K. A. Carlson, L. G. Harshman, Extended longevity lines of *Drosophila melanogaster*: abundance of yolk protein gene mRNA in fat body and ovary. Exp Gerontol 34, 173–184 (1999).

60. C. M. Gendron et al., *Drosophila* life span and physiology are modulated by sexual perception and reward. Science 343, 544–548 (2014).

61. Z. M. Harvanek et al., Perceptive costs of reproduction drive ageing and physiology in male *Drosophila*. Nat Ecol Evol 1, 152 (2017).

62. T. J. Maures et al., Males shorten the life span of *C. elegans* hermaphrodites via secreted compounds. Science 343, 541–544 (2014).

63. A. J. Spracklen, D. J. Kelpsch, X. Chen, C. N. Spracklen, T. L. Tootle, Prostaglandins temporally regulate cytoplasmic actin bundle formation during *Drosophila* oogenesis. Mol Biol Cell 25, 397–411 (2014).

64. S. Chen, fastp 1.0: An ultra-fast all-round tool for FASTQ data quality control and preprocessing. Imeta 4, e70078 (2025).

65. S. Chen, Y. Zhou, Y. Chen, J. Gu, fastp: an ultra-fast all-in-one FASTQ preprocessor. Bioinformatics 34, i884–i890 (2018).

66. A. Dobin et al., STAR: ultrafast universal RNA-seq aligner. Bioinformatics 29, 15–21 (2013).

67. Y. Liao, G. K. Smyth, W. Shi, featureCounts: an efficient general purpose program for assigning sequence reads to genomic features. Bioinformatics 30, 923–930 (2014).

68. M. I. Love, W. Huber, S. Anders, Moderated estimation of fold change and dispersion for RNA-seq data with DESeq2. Genome Biol 15, 550 (2014).

69. M. Ashburner et al., Gene ontology: tool for the unification of biology. The Gene Ontology Consortium. Nat Genet 25, 25–29 (2000).

70. G. Yu, Thirteen years of clusterProfiler. Innovation (Camb) 5, 100722 (2024).

